# What if I had a third arm? An EEG study of a supernumerary BCI system

**DOI:** 10.1101/817205

**Authors:** Jaime A. Riascos, David Steeven Villa, Anderson Maciel, Luciana Nedel, Dante Barone

**Affiliations:** Autonomous University Corporation of Nariño - Pasto, Colombia; SDAS - Research Group., Porto Alegre, Brazil; Institute of Informatics - Federal University Rio Grande do Sul (UFRGS) - Porto Alegre, Brazil; Ludwig-Maximilians-Universität München (LMU Munich) - Munich, Germany

**Keywords:** Motor Imagery, Brain-Computer Interface, Rubber-Hand Illusion, Embodied Cognition

## Abstract

Motor imagery Brain-Computer Interface (MI-BCI) enables bodyless communication by means of the imagination of body movements. Since its apparition, MI-BCI has been widely used in applications such as guiding a robotic prosthesis, or the navigation in games and virtual reality (VR) environments. Although psychological experiments, such as the Rubber Hand Illusion - RHI, suggest the human ability for creating body transfer illusions, MI-BCI only uses the imagination of real body parts as neurofeedback training and control commands. The present work studies and explores the inclusion of an imaginary third arm as a part of the control commands for MI-BCI systems. It also compares the effectiveness of using the conventional arrows and fixation cross as training step (*Graz* condition) against realistic human hands performing the corresponding tasks from a first-person perspective (*Hands* condition); both conditions wearing a VR headset. Ten healthy subjects participated in a two-session EEG experiment involving open-close hand tasks, including a third arm that comes out from the chest. The EEG analysis shows a strong power decrease in the sensory-motor areas for the third arm task in both training conditions. Such activity is significantly stronger for *Hands* than *Graz* condition, suggesting that the realistic scenario can reduce the abstractness of the third arm and improve the generation of motor imagery signals. The cognitive load is also assessed both by NASA-TLX and Task Load index.

## 1. Introduction

The primary purpose of Human-Computer Interaction (HCI) is seeking new alternatives for communicating humans and machines, and this effort is more evident when users with motor disabilities show difficulties using standard interfaces [1]. Brain-Computer Interface (BCI) is the technology that enables bodyless communication with machines or devices; this is done using the translation of brain signals into command outputs [2].

BCI commonly employs the electrical activity in the brain (EEG) elicited during a specific task. Depending on the nature of this activity, BCI is characterized as passive, active or reactive [3]. Passive systems use signals that arise without voluntary control. It is used fundamentally to asses mental states and enhance the human-computer interaction [4]. Active BCI works with the self-induced brain activity produced by the user independently of external events. It has been used as a control signal [5]. Finally, reactive BCI relies on the signals elicited by the reaction to specific external stimuli, which could be used to control an application as well [6].

Since the activation patterns of imaginary body movements involves both brain regions (sensory and motor areas) and neural mechanisms similar to the executed movement [7], the Motor Imagery BCI (MI-BCI) has been widely used and explored in active BCI [8]. MI-BCI employs the amplitude changes voluntarily elicited by the mental rehearsal of physical motor actions. Such variations are known as event-related de-synchronization and synchronization (ERD/ERS). These patterns have been successfully used for studying the neural mechanisms associated with motor actions, as well as a feature for classification in motor-related BCI systems [9, 8, 10, 1].

Despite BCI being a promising and useful application, there are still several challenges to be addressed. Chavarriaga et al. [11] discuss concrete research avenues and guidelines to overcome common pitfalls in BCI. Their paper is the outcome of a meeting held at the workshop “What’s wrong with us? Roadblocks and pitfalls in designing BCI applications”. They summarize four main topics that influence any closed-loop BCI system:

a. **Signal processing and decoding:** the signal processing of EEG data, and consequently BCI systems, is boosted by the fast growth of machine learning and unsupervised systems (i.e., deep learning) [12].
b. **End Users:** the creation of objective either questionnaires or pretests to identify potential user should be considered prior to a BCI implementation.
c. **Performance metrics and reporting:** BCI’s metrics are a topic under discussion [13] since the classification accuracy is not enough for evaluating BCI systems, the creation of new metrics becomes fundamental [14].
d. **Feedback and user training:** several efforts have been made in order to include the user inside the BCI loop [15], creating affordable and intuitive interfaces, considering human factors on their design.

In effect, immersive technologies have recently played an essential role in overcoming the feedback and user training challenge. Among them, Virtual Reality (VR) is one of the most promising technologies, giving the users a sensation of actual presence in virtual worlds. VR has been effectively used in several areas, from health-care for rehabilitation and training [16] up to data visualization and serious games [17, 18]. Likewise, VR has been used in BCI for a visual presentation feedback of the current task carried out by the user. Lécuyer et al. [19] discuss some of the current applications developed using BCI with VR, namely MindBalance [20], Simulation of wheelchair control [21], and “use the force” [22]. These studies, as highlighted by the authors, show the successful use of VR with BCI.

Another important thing about MI-BCI applications is that, so far, they have essentially used attached body parts. In other words, MI-BCI focuses on mental representations of jointed limbs following the human anatomy constraints (e.g., two arms, two legs, two feet, in a symmetrical distribution). To the authors’ knowledge, nevertheless, there are neither explorations nor applications that include non-embodied human limbs in BCI systems, even though Rubber Hand Illusion (RHI) experiments demonstrated the human capabilities to create body transfer illusions [23, 24]. Indeed, RHI does not only demonstrate a static body illusion representation (sense of ownership), but also an active movement eliciting a body illusion (sense of agency) [25].

In that vein, this paper presents the complementary results regarding the inclusion of a third arm as a control command in a BCI system: an EEG analysis of the induced brain oscillatory activity elicited by the third arm using Event-Related Spectral Perturbation (ERSP). A preliminary study addressed an offline exploration of the classification of the third arm task [26]. Continuing with that study, throughout this research, we compared the approach under two training conditions: the conventional Graz paradigm (cross and arrows) and immersive human-like feedback. Moreover, we included a cognitive load assessment by both the subjective questionnaire (NASA-TLX) [27] and the Task Load Index using EEG data [28]. Finally, we used the Movement Imagery Questionnaire −3 (MIQ – 3) [29] before the experiment to assess the movement imagery ability of the users. The findings suggest that ERS/ERD patterns are elicited by the virtual third arm. Moreover, in line with the literature, the realistic training enhances the modulation of such patterns but creating an additional cognitive load (presumably caused by the visual processing).

The remainder of this paper is structured as follows: section 2 presents the state-of-art in BCI, applications that use either VR or body illusions. Then, section 3 shows the materials, methods, and details of the experimental procedure. Finally, section 4 provides the main findings that are discussed in section 5, and section 6 presents the concluding remarks.

## 2. Related Works

Virtual Reality is a powerful tool to improve the BCI training and enhancing the feedback experiences [30]. The learning task should include an intuitive feedback so that the users can easily understand the action to be executed and improve their performance. However, it is currently hard to choose the right feedback presentation, and it should be a motivating and engaging environment [11], besides being natural and realistic. Here, VR can be shown as a real alternative for tackling the feedback presentation issue.

Lotte et al. [31] show how combining BCI with VR can carry towards a new and improved BCI system. Nevertheless, such VR feedback can also introduce some interference to the motor imagery-related brain activity used by the BCI because both *µ* and *β* bands are reactive in motor imagery and observation of the real movement [9]. An interesting study carried out by Neuper et al. [32] explores the influence of different types of visual feedback in the modulation of the EEG signal during the BCI control. Using a video to show a first-person view of an object-directed grasping movement, they were able to found modulation activity in sensorimotor rhythms caused by this real feedback stimulus. They highlight the importance of the amount of information provided by this condition in order to reduce the reactive bands.

Ron-Angevin and Diaz-Estrella [33] made a first comparison between the screen condition (Graz) and VR in a BCI scenario, focusing on the performance (classification rates). They successfully found improvements in the feedback control of the VR condition in untrained subjects. However, they used car navigation as a task, which can be seen as unnatural and abstract when compared to an embodied experience. The studies cited above have used different feedback stimuli, but none of them has used a virtual human avatar, which could be useful for the training step. Recently, Skola and Liarnokapis [34] addressed such problem comparing the Graz paradigm against a human-like avatar performing the user’s motor actions synchronously. The authors report improvements in both ERD/ERS modulation and classification rates by the neurofeedback-guided motor imagery training. Likewise, Braun et al. [35] report the same sort of results using an anthropomorphic robotic hand as a visual guide. Also, they found differences between the two conditions in the electrodermal activity and subjective measures. Both works reported that they were inspired by the RHI. They also include within their discussions, the analysis of sense of ownership, agency, and self-location towards the non-body object, concepts that are being recently taken into account in BCI research [36, 37].

Although, from the RHI theory, it is demonstrated that the body transfer illusion can be effectively used with non-attached limbs in both passive (presence) and active (movement) conditions [25]. Up to this point, supernumerary limbs BCI system had not been approached. Bashford and Mehring [38] proposed this possibility with their work. They used an imaginary third arm for assessing the ownership and agency of a non-body limb in an imitation BCI (i.e. subjects think that their EEG activity is controlling the arm). Results show that there is independent ownership and control – based on the correct movements observed against the subject movements – of the third arm keeping the sense of ownership of the real hands. These findings suggest the capabilities of human of extrapolating limbs to execute motor actions. However, they did not study the use of this third arm as a control command inside the BCI loop. A recent work proposed by Song and King [39] demon-strates that using an RHI-based paradigm can significantly enhance the MI signals for BCI systems.

The paper’s contribution includes a step towards the creation of supernumerary MI-BCI systems. Here, we performed an EEG study of the user’s ability to imagine a third imaginary arm in a BCI paradigm. We also compared the effectiveness of using the conventional arrows and fixation cross as training step (*Graz*) against a first-person view using a human avatar (*Hands*). Both training conditions were carried out in a VR environment.

## 3. Materials and Methods

### 3.1. Overview

An offline MI-BCI experiment, which uses EEG for recording the data and VR scenarios for presenting the stimulus, was conducted in a reduced noise room. The experiment’s aim is to study the feasibility of including a virtual third arm in a MI-BCI system while the traditional training paradigm (Graz) is compared against a first-person view using a human avatar. There were two recording sessions with two runs in each one with a resting time between them. The sessions were conducted on two separate days within one week. Only on the first day, the participants had to fill up three questionnaires: MIQ-3, demographics and Edinburgh Handedness. Likewise, after each session, participants filled the NASA-TLX form.

### 3.2. Participants

Ten right-handed volunteers (four women) participated in the study. Participant ages were within 18 and 34 years old with a mean of 23. All participants had basic informatics knowledge. Only 30% did not have previous experience with VR and no one had any previous experience in MI-BCI. No one had problems with head movements. Half of the population had visual impairments (mainly myopia and astigmatism) and used glasses to reduce them. The experiment was conducted in accordance with the Declaration of Helsinki. Participants were informed both oral and writerly about the procedure and the EEG recording. All participants gave written informed consent.

### 3.3. VR Scenarios

For the VR exposition, we used a head-mounted display (HMD) Oculus Rift CV1 with a resolution of 2160 × 1200 (1080 × 1200 per eye), refresh rate of 90 Hz, a 110° field of view, and both rotational and positional tracking to render the immersive scene. We used the popular game engine Unity3D to develop the immersive scene that was intended to assist the users when imagining and performing motor actions with their left and right real arms and the middle imaginary one (see the top of Figure 1).

**Figure 1:**
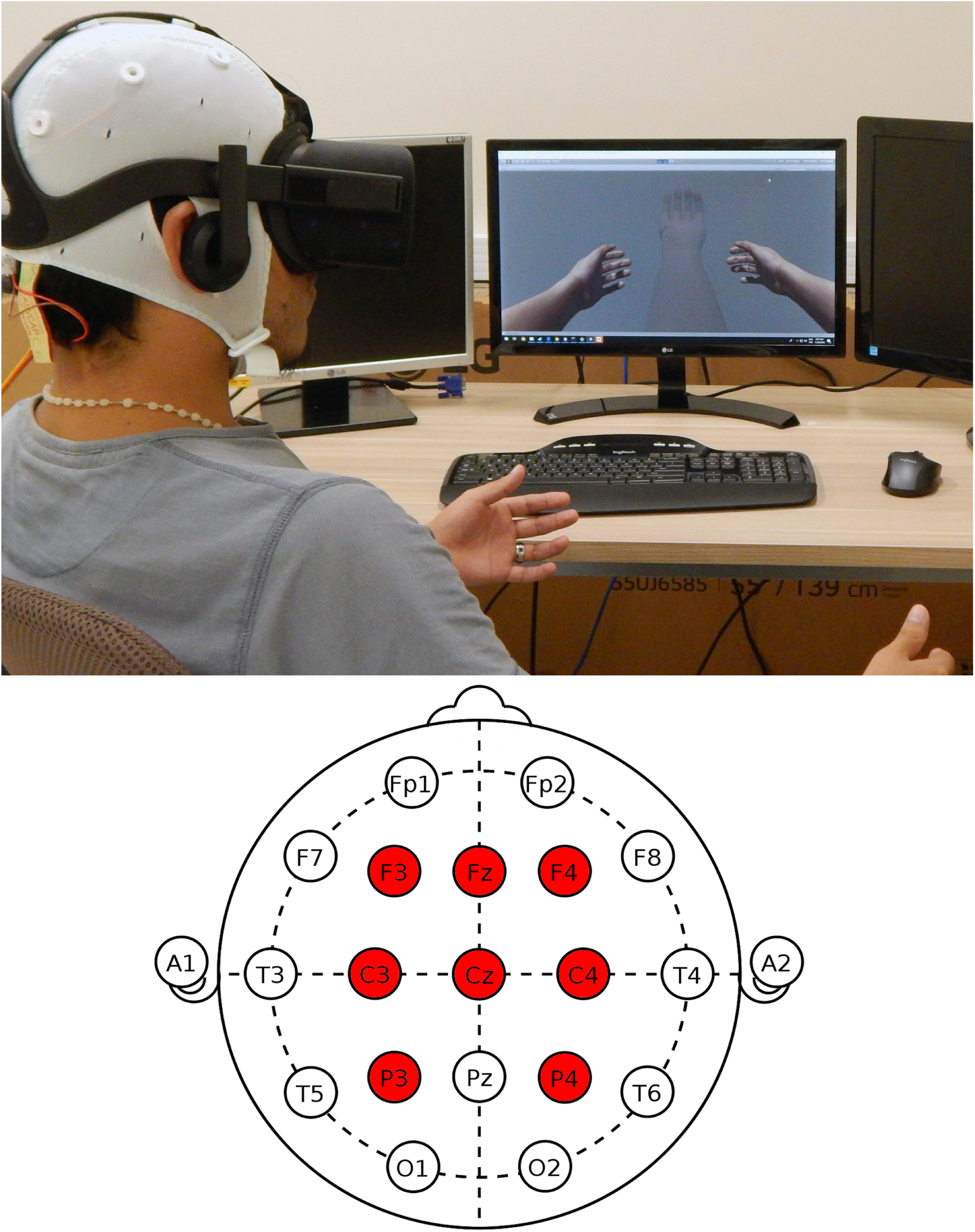
Experiment setup: A subject using a BCI interface to control his “three” arms in a virtual reality experience (top); and the electrodes placement over the sensorimotor area (filled circle), following the 10-20 system (bottom).

There was a special focus on the realism of the models: left and right hands were placed matching with the rest positions of the real hands. A third hand was placed in the middle of the body, like emerging from the chest trying to avoid visual relations with the left or the right arm. The fingers on the third arm also were modified to be symmetric. In this sense, since that the thumbs in either left and right hand can indicate to which arm it belongs, their were removed from the third arm. Thus, it is identified as an independent arm and not a copy or extension of the existing arms. High-quality textures were used with shaders designed to highlight generic skin details. Bones in each finger preserve the average human hand proportions.

### 3.4. Experimental Procedure

The subjects sat comfortably in an armchair and were asked to rest their arms in the armrests and avoid any other movements during the recordings. Initially, the participants wore the HMD for getting into the scene and running several trials for learning the instructions previously read. After the training, we mounted the EEG cap followed by the traditional gelling process, and then we fit the HMD. We tried as much as possible to avoid that the HMD frame touches the EEG electrodes. Moreover, we checked the signal quality before and after mounting the HMD to detect any avoidable interference.

The experiment involves the execution of four different tasks in two experimental conditions. The subjects were invited to rest (RS), or to move a specific hand: third hand (TH), left hand (LH), and right hand (RH). Conditions considered were *Graz*, and *Hands*. The *Hands* condition involved the presentation of a human-like avatar (see the top of Figure 2), whereas *Graz* the presentation of arrows (see the middle of Figure 2).

**Figure 2:**
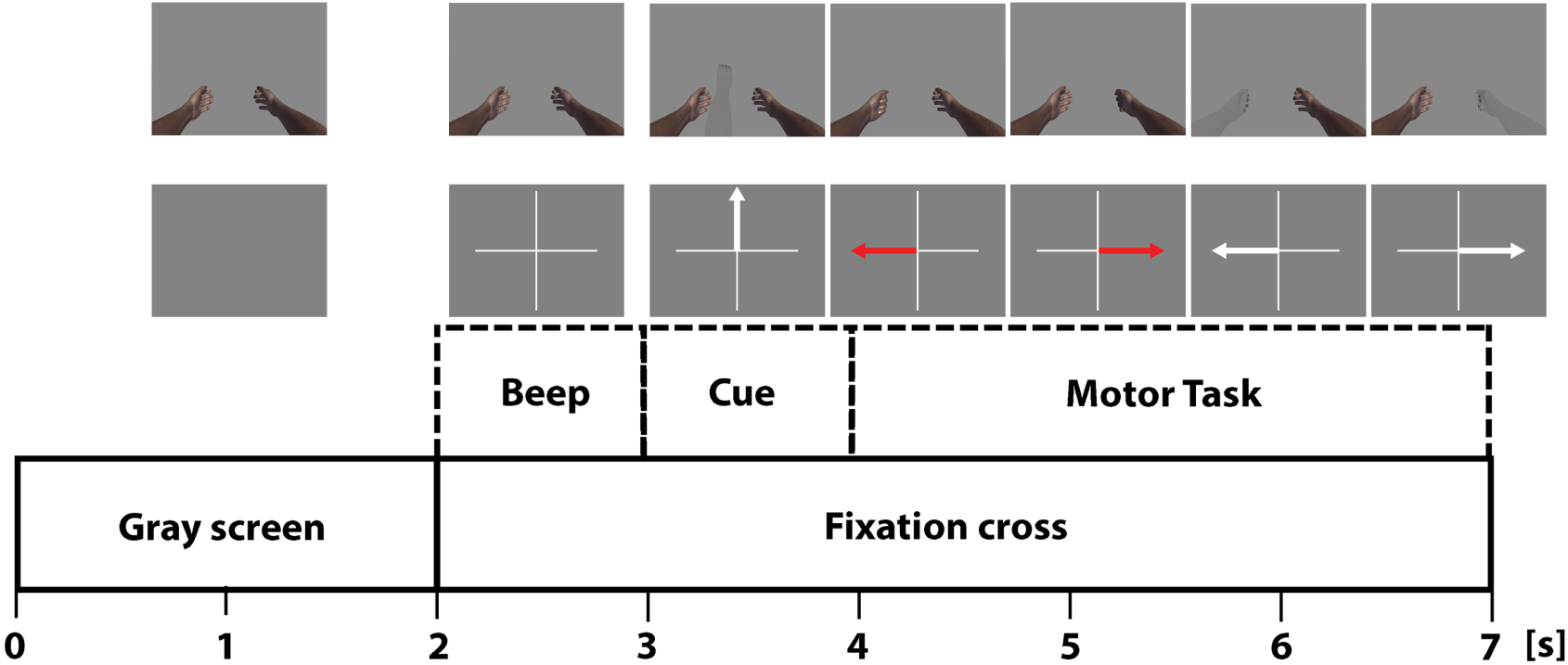
Experiment paradigm. The visual stimulus of the task’s cue are corresponding for both conditions. Top: visual stimuli for *Hands* condition. Middle: visual stimuli for *Graz* condition. Bottom: timing of the trials following the classic Graz protocol.

The two experimental conditions followed the timing protocol proposed by Pfurtscheller [9]. The users performed 20 trials of each task randomly selected (described below) with a duration of 7 seconds each (see the bottom of Figure 2). The main difference between the conditions lies in the visual feedback, as follows:

a. ***Graz* condition**: starting with a gray screen (resting state), at time 2s, a fixation cross at the center of the scene was displayed with a short warning tone (‘beep’) which indicates to the user to pay attention to the incoming visual cue presented at time 3s. At time 4s, the user had to perform the motor task for three seconds. The color of the arrows indicates the task (red for execution and white for imagination) and the direction indicates if the hand should be either left or right. The third arm cue was an arrow pointing upwards (see the middle of Figure 2).
b. ***Hands* condition**: at the start, the user’s hands were placed in the equivalent real arms positions (resting state), at time 2s, the same auditory cue starts indicating an incoming stimulus. Next at time 3s, a visual cue is introduced without animation to let the users to be prepared for the action they will perform. At time 4s, the animation is introduced, and the user must perform either the mechanic or imaginary operation. This state continues until the end of the task (three seconds more). As for the visual cues, the real skin shading represents actual open-close hand movements, while transparent shading represents imaginary movements. Moreover, it is important to highlight that the third arm appears in the scene only when this specific trial is necessary. In other trials, there are just two visible hands (see the top of Figure 2).

Following [40], subjects were instructed to perform the kinesthetic experience during the execution of motor imagery tasks, i.e., imagining the sensation of performing the motor tasks rather than the visual representation of the movement. The authors suggest that kinesthetic motor imagery is essential to elicit sensorimotor patterns (ERD*\*S). Besides this, in order to avoid the carry-over bias, both experimental conditions were counterbalanced across participants (i.e. five subjects start with *Hands* condition and the rest with *Graz*). Likewise, it is necessary to mention that the movement animations were applied directly to the bones always looking for a natural behavior of the hand. The animations are predefined, they are not based on the user’s EEG activity or motion.

Finally, in contrast to Skola and Liarnokapis [34] where the *Graz* condition is presented in a monitor, we made comparisons of the *Graz* and *Hands* conditions in an immersive virtual environment. Therefore, the users have to wear the HMD in both conditions. The background of Graz scenario was set to gray, avoiding high contrast that could produce discomfort on the user’s eyes.

### 3.5. Data Acquisition

We collected the EEG data using an OpenBCI 32 bit board at a sampling rate of 250 Hz. Following the 10-20 EEG placement system, eight passive gold cup electrodes were used and placed at sensorimotor cortex (see the bottom of Figure 1), namely, frontal (F3, Fz, F4) central (C3, Cz, C4), and parietal (P4, P3) cortices. Left and right mastoids were used as reference and ground electrodes respectively. Labstreaminglayer (LSL) is used for recording and synchronizing the EEG data with the Unity trials through LSL4Unity (a third party software) [41].

### 3.6. EEG signal processing

We used EEGLAB (14.1) [42] (under Matlab 2017b) for processing the .XDF file created by LSL. Following the usual procedure for analysis motor-imagery-related EEG patterns (sensorimotor rhythms) [43], we initially down-sampled the signals at 115 Hz and band-passed at 1-35Hz using a finite impulse response (FIR) filter. Later, we used the Cleanline plugin at 50-115 Hz instead of a notch filter to avoid band-holes, and distortions at the cutoff frequency. Likewise, we rejected bad channels (excluding the sensorimotor ones) using Cleanraw plugin. The rejected channels were then interpolated using a spherical function. Finally, we used the common average reference (CAR).

### 3.7. Event-related spectral perturbation

The event-related spectral perturbation (ERSP) is a generalization of the ERD/ERS patterns. ERSP computes the changes of the spectral powers in time-frequency domains, relative to the stimuli [44]. Thus, with this approach, the changes of the EEG signals elicited by motor imagery events can be detected alongside of the spectral band and epoch. ERSP values were computed for every mental task (TH, LH, RH, RS) in *Graz* and *Hands* conditions using the *newtime* function of the toolbox in the filtered data. We used a time window of −500 ms to 2500 ms, displayed between 5 Hz and 30 Hz; Also, significant alpha was setup to 0.05. The sensorimotor area composed by the electrodes C3, Cz and C4 were used to display the time-frequency ERD/ERS maps (Figures 3 and 4).

**Figure 3:**
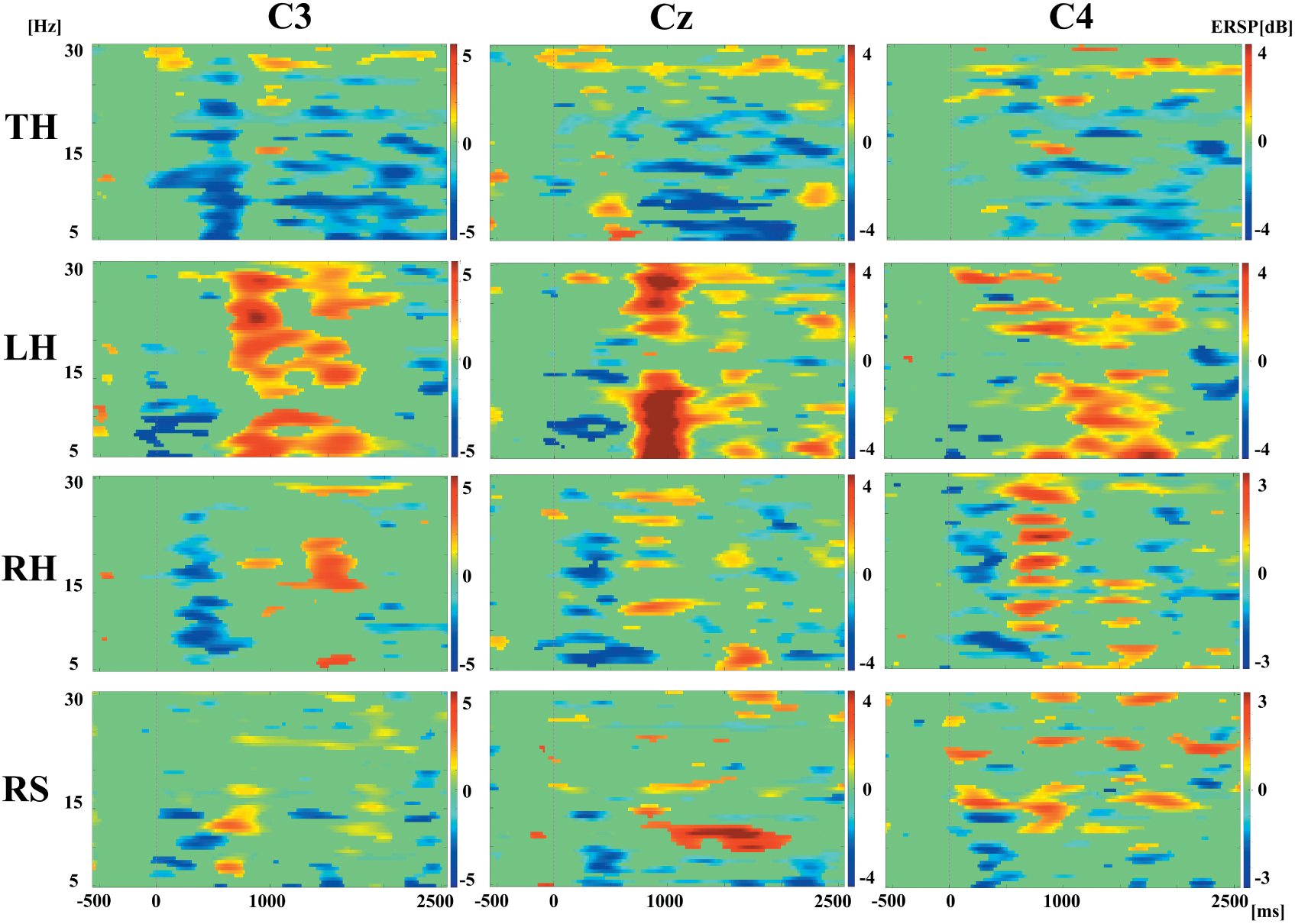
Significant ERD/ERS patterns of the mental task at C3, Cz, C4 positions for *Hands* condition (blue indicates ERD). A strong ERD activity is found at the three electrodes for the third hand (TH). Whereas, ERS patterns are found mainly for the left hand (LH). The ERD/ERS fluctuation is more visible for the right hand (RH), mostly at C4.

**Figure 4:**
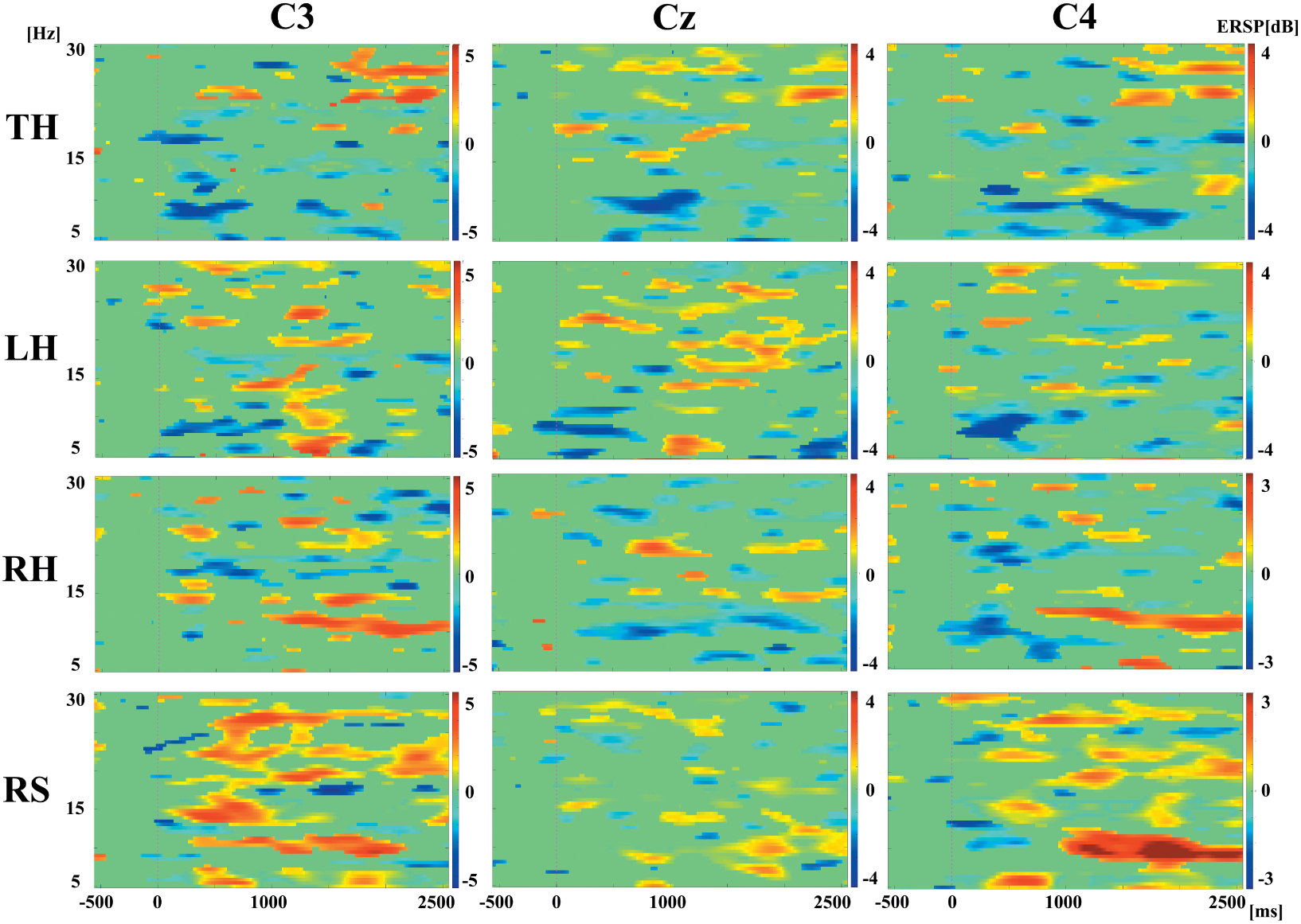
Significant ERD/ERS patterns of the mental task at C3, Cz, C4 positions for *Graz* condition (blue indicates ERD). An ERD activity is mainly found in the alpha band (8-12 Hz) at the three electrodes for the third hand (TH). The ERD/ERS patterns are widespread for left and right hands (LH, RH respectively) at the three electrodes. There is extensive activity in the resting state (RS).

### 3.8. Task Load Index

Besides the subjective assessment of the cognitive load by the NASA-TLX [27], we also used the Task Load Index (*TLI*) developed by Alan Gevins and Michael E. Smith [28] in order to have an objective measure of the task load. The authors found that the power changes of *θ* at frontal mid-line sites and *α* at parietal sites are related to the task load associated to the mental effort required for task performance. Thus, this index can be measured by the ratio of *θ* to *α*. In this work, we used the *spectopo* function to calculate the average of the absolute power of frontal mid-line (F3, Fz, F4) *θ* and parietal (P3-P4 plus Cz) *α* to assess the mental tasks per condition (*Graz* and *Hands*) as follows:

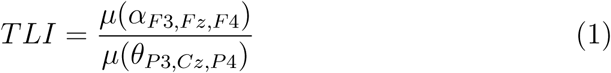

### 3.9. MIQ-3

Despite Motor Imagery is a fundamental constructor of any healthy person, i.e., that all humans should have the capacity of imagining and planning motor activities, some people could face limitations to perform imaginary activities. In such vein, several questionnaires were made in order to subjectively assess the individual ability to perform imaginary motor tasks, such as the Vividness of Movement Imagery Questionnaire (VMIQ) [45] or Movement Imagery Questionnaire (MIQ) [46]. The MIQ-3, a recent version of the MIQ, and in different of the VMIQ, assesses three kinds of imagery [47]:

a. **Internal Visual Imagery**: visual image of the performed movement from an internal perspective (i.e., the subject performing and seeing the action from a 1st person perspective).
b. **External Visual Imagery**: visual image of the performed movement from an external perspective (i.e., the subject performing and seeing the action from a from a 3rd person perspective)
c. **Kinesthetic Imagery**: creating the feeling of making the performed movement without actually doing it.

This survey is a 12-item questionnaire to asses the capacity to image four simple movements: a knee lift, jump, arm movement, and waist bend, in a scale from 1 (very hard) to 7 (very easy). The MIQ-3 demonstrated excellent psychometric properties, internal reliability, and predictive validity. This paper uses an adaptation of the MIQ-3 questionnaire to the Portuguese language [48].

## 4. Results

### 4.1. ERSP maps

Figures 3 and 4 show the time-frequency representation of significant (bootstrap method, *p <* 0.05) ERD/ERS values (blue indicates ERD) for the *Hands* and *Graz* condition respectively. These maps come from a single subject (*6*)^1^ at electrode positions C3, Cz, and C4. The analysis of these maps reveals, certainly, the brain activity elicited by the imagination of hands movements (motor imagery), and that the third arm emerging from the chest can elicit similar patterns.

For the TH task, at C3 position in *Hands* condition, a strong power decrease is clearly visible around 500ms after stimulus onset, and this behavior repeats in almost the whole frequency range. In the other two imagery tasks, LH has a decrease in Alpha followed by an increase in Alpha and Beta. RH has a similar pattern but without a clear ERS activity in alpha. Interestingly, TH task held the ERD activity during the rest of the epoch after 1000ms with few ERS in middle and high beta bands. Conversely, in Graz condition at C3, the ERD patterns of the TH task are attenuated and widespread with some ERS activity at the end of the epoch in high Beta band.

At Cz in *Hands* condition, the TH task presents a few ERS activity that starts around 500ms in Alpha, and an ERD that starts around 1000ms in Alpha and Beta bands. LH presents a strong ERS activity in both Alpha and Beta anticipated by an ERS in Alpha and middle Beta. RH has a strong ERD activity in Alpha and Beta and posteriorly some ERS in high and low Beta. Meanwhile, in *Graz* condition, TH shows ERD patterns in Alpha until the first 1000ms. At the end of the epoch, some ERS activity is presented in high Beta. In LH, there is an ERD pattern in Alpha during the first 500ms and a widespread ERS activity later. RH holds the ERD in Alpha at the same time with some ERS in middle Beta.

Similarly, TH task in *Hands* condition presents an ERD pattern around 500ms in Alpha and middle Beta at the C4 position. This activity is held again during the whole epoch (mainly in Alpha). Few ERS activity is found in high Beta after 1000ms. The ERS activity is most prominent in Alpha and low-middle Beta for LH, meanwhile, RH shows an ERD/ERS pattern in Alpha and Beta in the first 1000ms. For *Graz* condition, the ERD patterns of TH task are widespread in Alpha and Beta between 500ms and 1500ms with some presence of ERS in high Beta. LH has a strong ERD activity during the first 1000ms in Alpha and some widespread ERS in high Beta. RH has strong ERD patterns during the same previous time in both Alpha and middle Beta followed by a strong ERS activity in Alpha, extended along of the epoch.

### 4.2. Topographical Maps

Figure 5 shows the representative set of topographical distributions of each mental task obtained from the same subject in Alpha and Beta bands for the first second after the cue. The TH task, for both bands in *Hands* condition only exhibits ERD activity (more prominent in Alpha band) mainly on the contralateral (C3) and middle (Cz) regions. On other hand, TH in *Graz* condition presents ERD/ERS activity in both bands; in effect, it can be seen a strong ERD on the frontal lobe (F3) and ERS on parietal region (P3). These findings could suggest that the brain activity elicited by the third arm is not only associated with sensorimotor areas, but also the imagination effort is visible at frontal and parietal regions (more clear in *Hands* condition for both bands).

**Figure 5:**
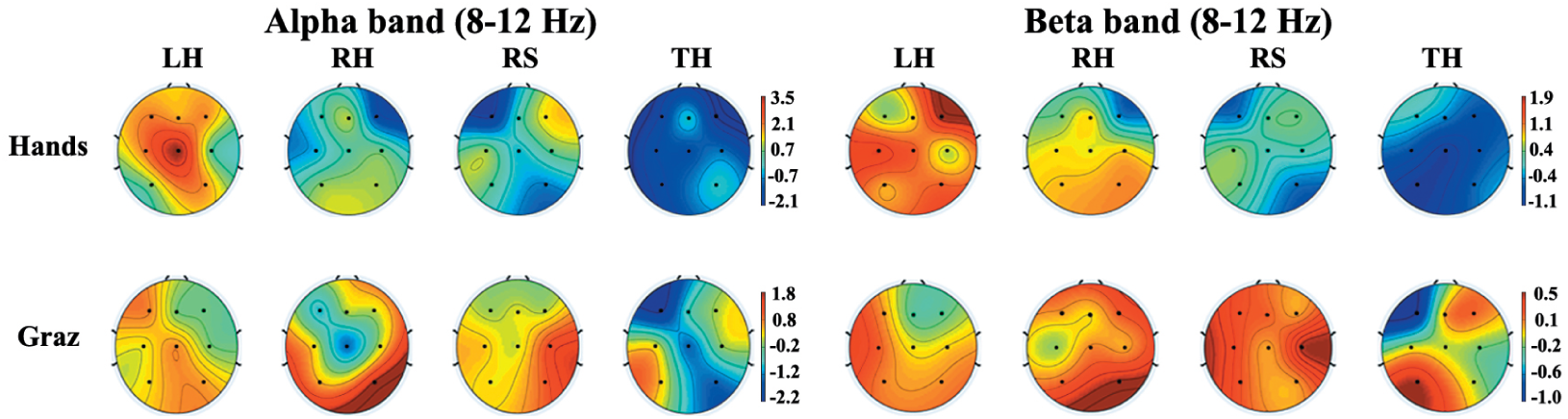
Topographical distribution of each task for both conditions (blue indicates ERD). The maps are made using the ERSP values in both Alpha and Beta bands, one second after the cue. Blue indicates the ERD activity during the mental tasks.

### 4.3. Power spectral analysis

In order to explore the differences of the ERD/ERS patterns among tasks in the two conditions, Figures 6 and 7 show comparisons of the power changes of the TH task against the other imagery tasks (LH-RH) in both conditions using the same electrodes array from the same subject (*6*). Blue lines represent TH, the red ones LH while RH is represented by green lines. Moreover, Figure 8 presents the power comparison of the TH task in both conditions. Blue line indaicates The paired Wilcoxon signed-rank test was used to find out significant differences between conditions (*p <* 0.05). They are indicated by shaded blocks.

**Figure 6:**
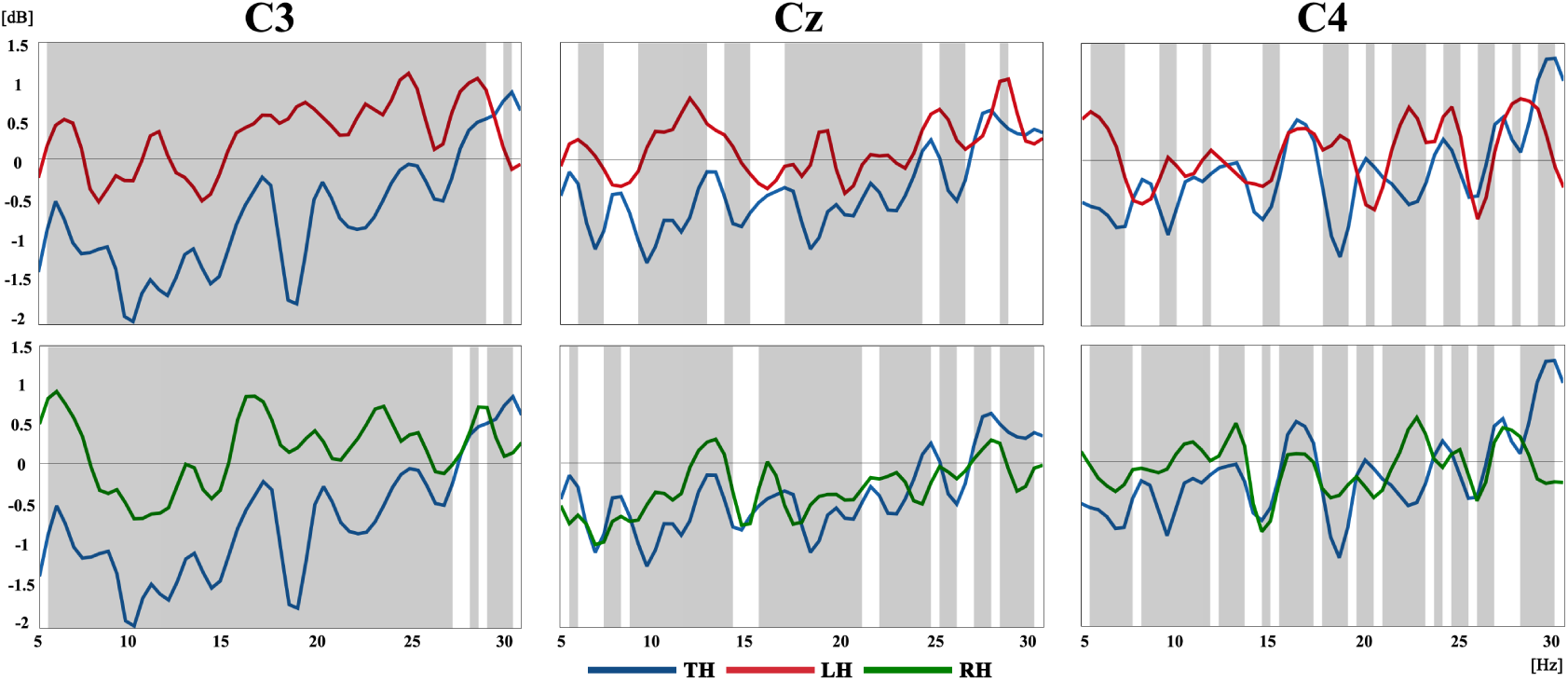
Comparison of the power changes of the mental tasks in the sensory-motor area (C3, Cz, C4) in *Hands* condition. Top: Third hand (TH, blue) - Left hand (LH, red). Bottom: Third hand (TH, blue) - Right hand (RH, red).

**Figure 7:**
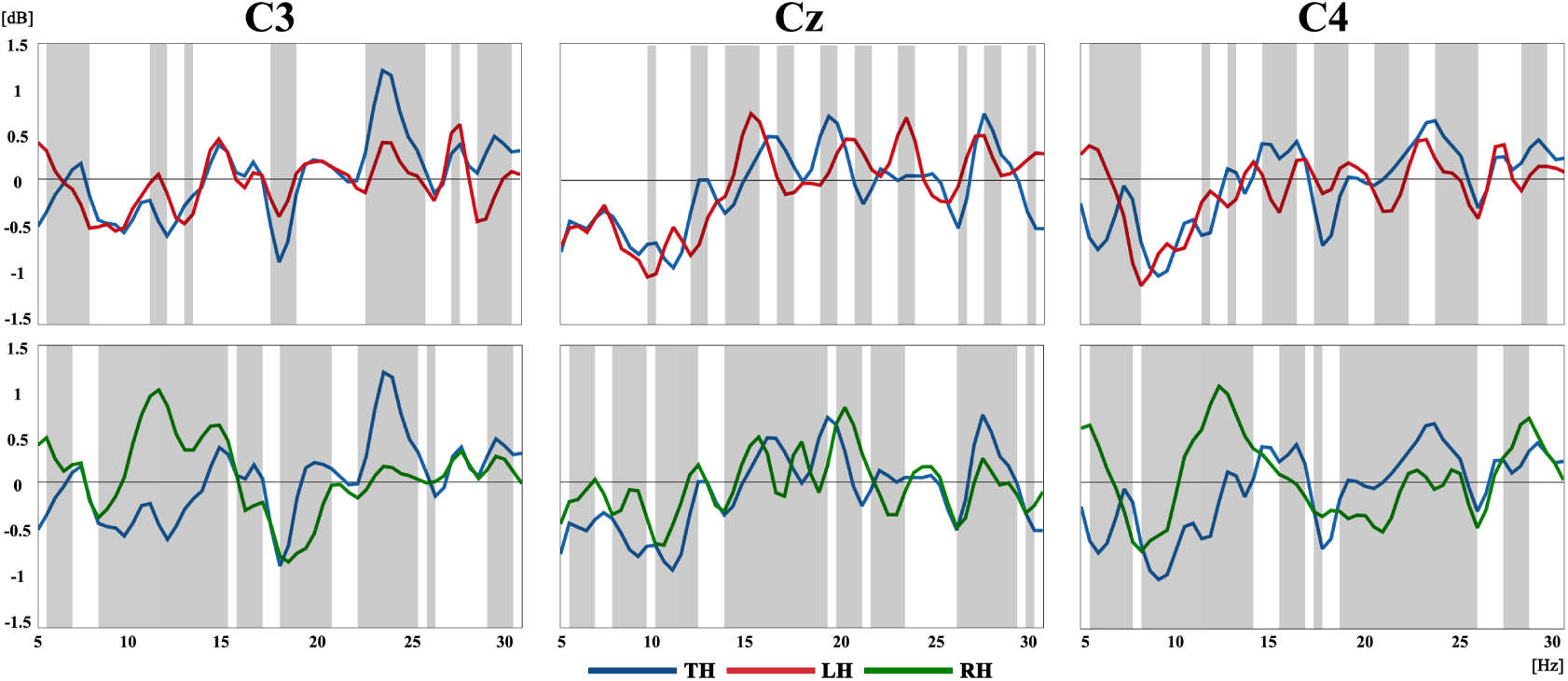
Comparison of the power changes of the mental tasks in the sensory-motor area (C3, Cz, C4) in *Graz* condition. Top: Third hand (TH, blue) - Left hand (LH, red). Bottom: Third hand (TH, blue) - Right hand (RH, red).

**Figure 8:**
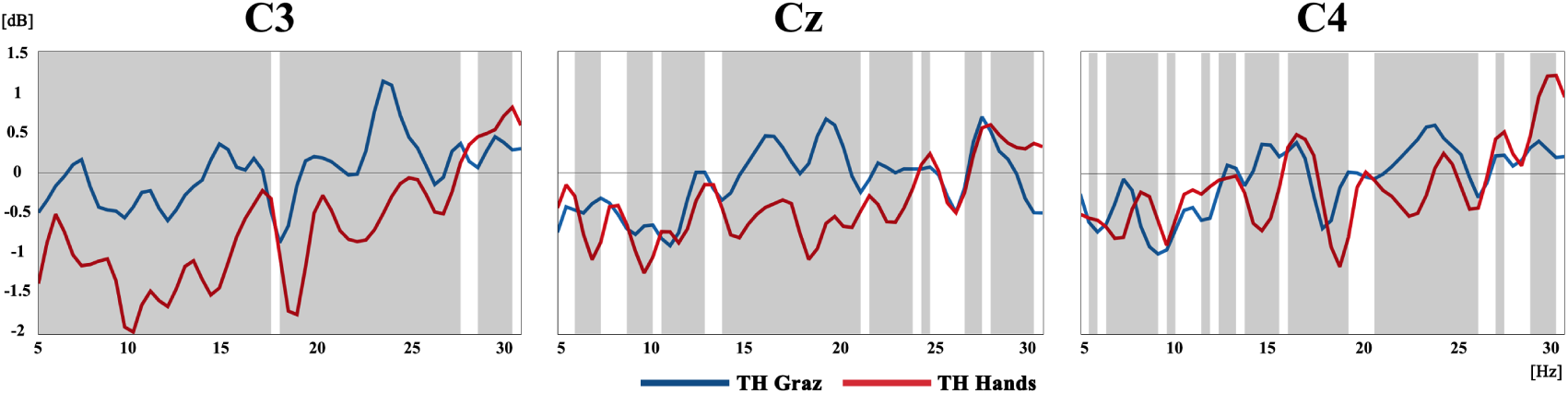
Comparison of the power changes of the third arm task in the sensory-motor area (C3, Cz, C4) for both conditions. Blue: Third hand in *Graz* condition. Red: Third hand in *Hands* condition.

The differences presented by TH-LH and TH-RH are significantly more broad-banded at C3 than other channels in *Hands* condition (Figure 6). Meanwhile, *Graz* condition presents similar significant region sizes among the channels (Figure 7). At C3, both cases (TH-RH, TH-LH) in *Hands* condition show significant differences in almost the whole frequency range. Conversely, in *Graz* condition, TH-RH shows more significant differences in Alpha and low Beta than TH-LH, but they share the significant region around 20Hz up to 25Hz.

At Cz in *Hands* condition, the TH-LH comparison does not have a significant region in the Alpha band, but it shares a low and middle Beta with TH-RH, which has significant differences in Alpha and high Beta sub-bands. For the *Graz* condition in the same location, the TH-LH comparison indicates wide-spread sub-band regions for the Beta, in Alpha only a small region around 10hz is presented and, in the meantime, TH-RH shows a consistent region in Alpha and low and high Beta.

Finally, at C4 in Hands, the TH-RH comparison shows wider regions than TH-LH, especially in Alpha and middle Beta rhythms. The same behavior is presented in the *Graz* condition, where TH-RH has more significant regions in Alpha and low and middle Beta than TH-LH, which does not have a significant difference in Alpha, only in several sub-bands along Beta, mainly above than 15Hz.

In the comparison of the TH task between conditions (Figure 8), there is a stronger power decrease in *Hands* than in *Graz* condition, in line with the ERS/ERD maps (Figures 3 and 4). Such difference is more evident at C3 than the other channels. Likewise, C3 noticeably shows significant regions within both Alpha and Beta rhythms, whereas Cz is more often in middle and high beta, and C4 in Alpha and middle Beta.

### 4.4. Cognitive Load and MIQ results

Figure 9 shows the cognitive load of both objective (Task Load Index) and subjective (NASA-TLX) analyzes. The results from the cognitive load assessed by the Task Load Index show that the *Hands* condition has a significantly higher cognitive load than the *Graz* one (pairwise paired Wilcox with Bonferroni: V = 656, p-value = 0.00063). There is no significant difference among the imaginary tasks (TH,RH, LH) and resting state (RS). Meanwhile, the subjective assessment of the cognitive load reflects the opposite. NASA Workload points to a higher cognitive load in the *Graz* condition instead, although significance could not be found (paired t-test: t=0.829, p-value=0.428).

**Figure 9:**
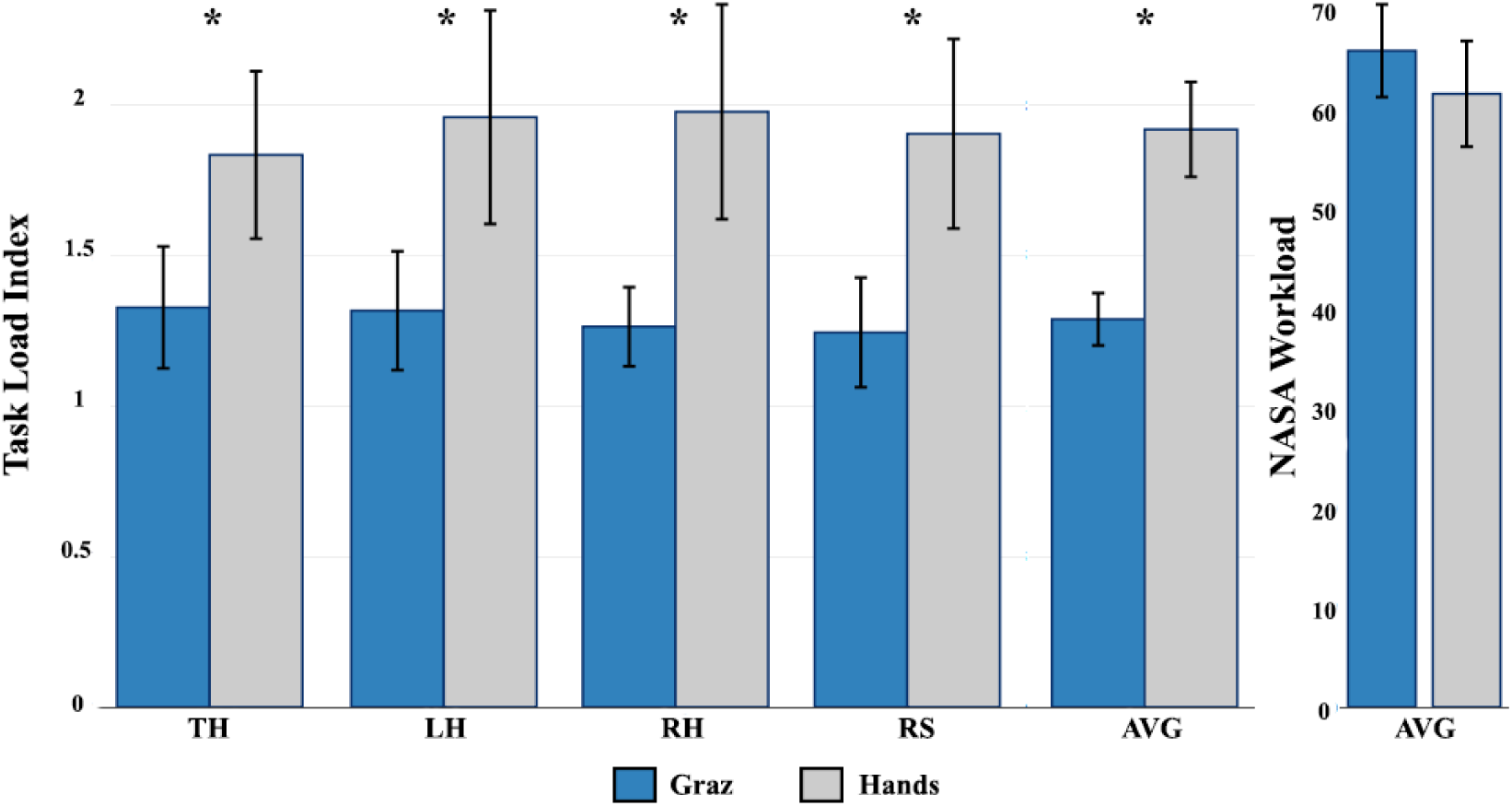
Task Load Index and NASA Workload assessment for the two conditions. * Significant differences

Figure 10 shows that the *Hands* condition presents a non-significant higher Load Magnitude than *Graz* in factors such as Performance, Physical and Temporal demand. Nevertheless, a pairwise paired Wilcoxon reflects that there is a significant difference between conditions in the Frustration factor (V=210, p-value= 0.049), indicating a higher sense of frustration in *Graz* than *Hands* condition.

**Figure 10:**
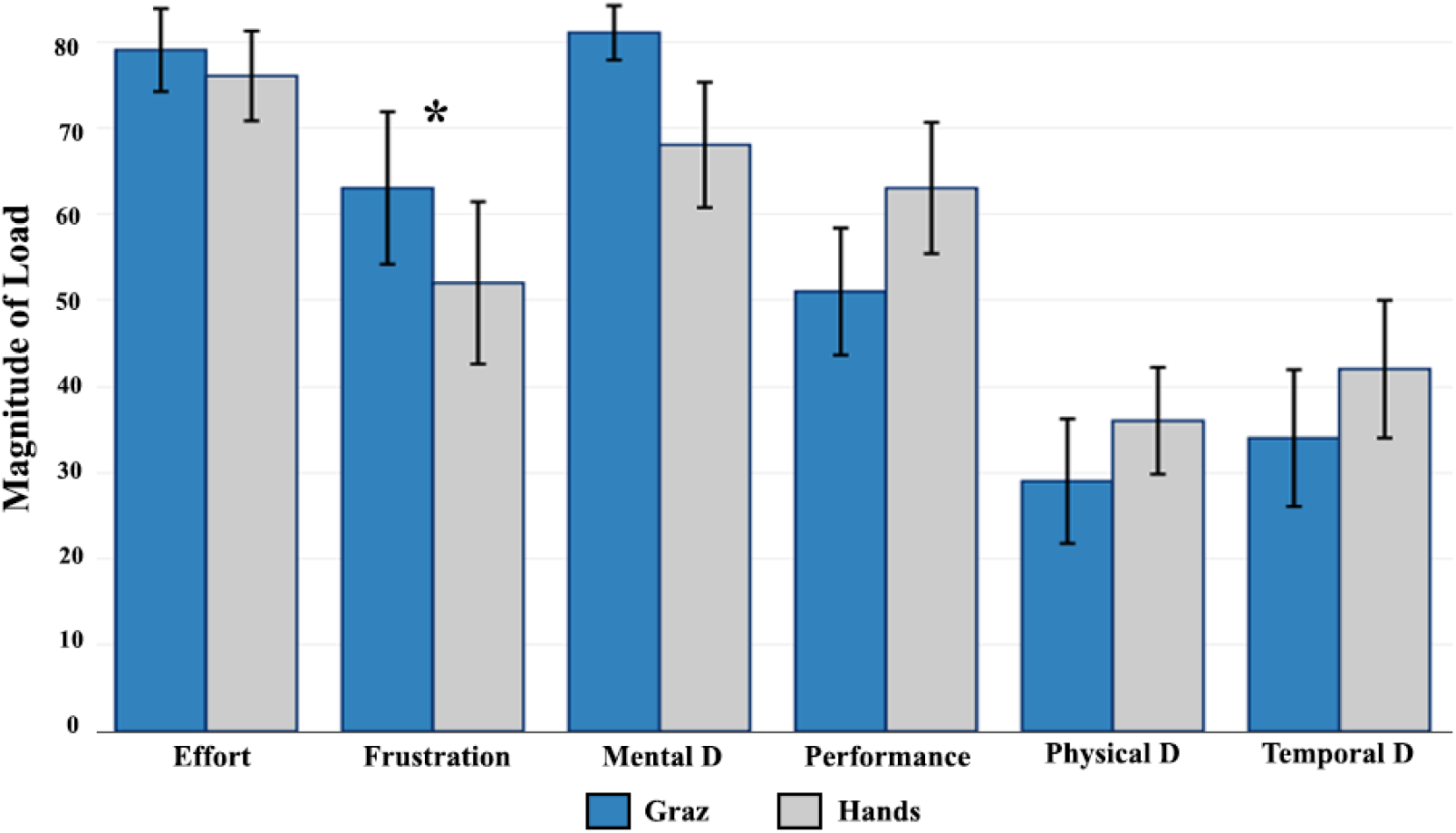
NASA factors for the two conditions. *Significant difference.

Finally, a study about the difficulty of performing imaginary tasks was carried out through the Mental Imaginary Questionnaire (MIQ-3). Figure 11 summarizes the user’s answers of the MIQ-3 questionnaire, the ratings represent how easy (7) or hard (1) was to perform the imagery task. The mean values show that External Visual Imagery (5 *±* 1.02) was easier for the users than Internal Visual Imagery (4.8 *±* 1.13) and Kinesthetic Imagery (3.95 *±* 1.24).

**Figure 11:**
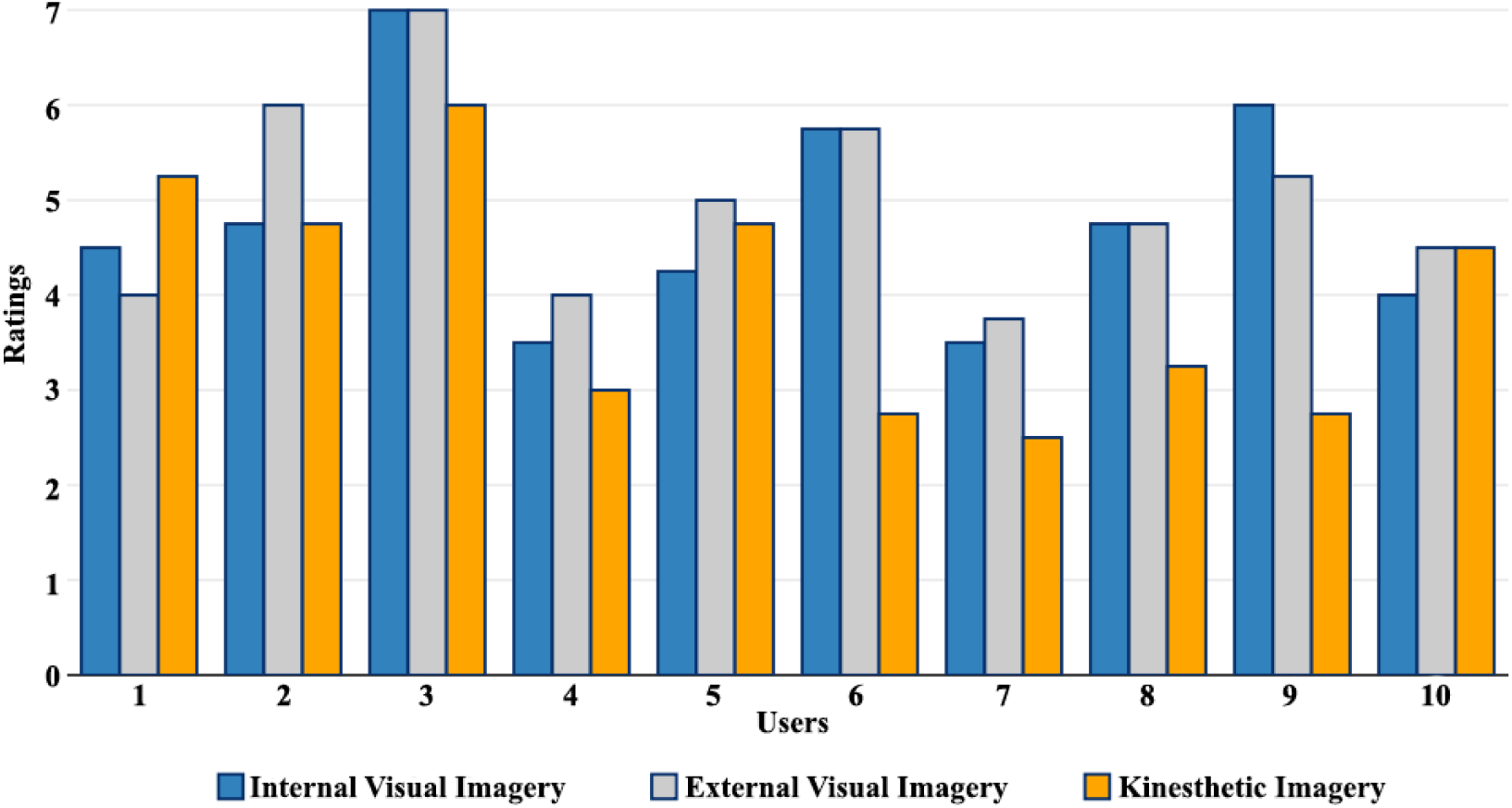
MIQ-3 results. Ratings range from 1 (very hard) to 7 (very easy).

## 5. Discussion

This study proposed the inclusion of a third arm in an MI-BCI application creating thus a supernumerary limb MI-BCI system. Furthermore, for this approach, the influence of embodiment feedback (*Hands*) was compared with the standard Graz training in VR. In line with the previous works [34, 35, 26], both the classification rates and the modulation of ERD/ERS signals were enhanced by the realistic feedback, evidencing its importance inside the BCI loop. Also, our work goes further than the one done by Skola and Liarnokapis [34] because they compared an embodied VR scenario against a monitor-based Graz, creating a bias in the users who started with VR. Here, the comparison was made with both Graz and Hands experimental conditions performed in immersive VR.

The presented patterns (Figures 3 and 4) suggest a significantly decreased activity in the sensorimotor area caused by the realistic feedback in comparison with the conventional paradigm (*Graz*). Besides this, the ERD activity of TH task is prominent at the three sensorimotor channels (C3, Cz C4) which could suggest that there is not a compulsory hemisphere governing the control and action of the imaginary third arm. Nevertheless, the analysis of the power changes between tasks (Figures 6 and 7) shows that there are more significant regions at C3 than at the other electrode positions. This result could indicate that the user’s handedness influences the region where TH task presents more activity. In the same way, the common ERD/ERS pattern is visible in LH and RH tasks, more in RH than LH; but it was missing in TH (only an increasing power activity was found in higher frequencies: > 25*Hz*). It could suggest that the absence of symmetry of the third arm does not elicit a supplementary ERS activity for this task, and this fact is visible in the topographical maps (Figure 5) where the TH in *Hands* condition presents only ERD activity. This could indicate an effect of the virtual arms support the users to create the abstraction of the third arm. Moreover, the unexpected activities presented in the resting state (RS) could be caused by the inertia of the execution/imagery movements. The paradigm to be adopted in the future should include a blank space between the motor task and resting state so that the movements could be easily excluded.

The aim of studying the cognitive load in both subjective and objective ways is for a deeper understanding of the additional load that realistic and visual feedback could cause. In effect, the outcome of the objective assessment (Task Load Index) is not supported by the results of the subjective one (NASA-TLX). EEG data reveals that the cognitive load is higher (significantly) in the realistic condition (*Hands*) than the standard one (*Graz*) but the opposite seems to occur in the NASA-TLX (without significance). Moreover, some user’s comments at the end of the experiment, such as “I found harder the arrows than the arms” or “I feel Temporal demand a bit easier in Hands than Graz because it is easier to visualize” and the opposite “… The arrow session was a easier than the virtual hands because with the arms I constantly tried to follow the hand movements which did not happen with the arrows” could evidence the disjunctive sensation of the users evidenced by the NASA and Task Load Index. Interestingly, a user did the next comment “The fact that I had the possibility of performing real hand movement helped me to release the stress created by the imagery tasks.” This comment supports our decision of keeping the real movements alongside of the imaginary ones, but further studies and comparisons are necessary before drawing conclusions. Finally, the imagery questionnaire shows that the External Visual Imagery was more natural to the users, complementing the comments of the users.

## 6. Conclusion and Future work

This study investigated the possibility of using an imaginary third arm in a BCI system, and shows the differences of the EEG patterns of using a realistic visual training in comparison of the traditional visualization. Initially, the common EEG patterns of motor imagery activity (ERD/ERS) are found when the subjects were asked to imagine a hand movement of a third arm emerging from the chest. These findings can suggest that the illusion of having a third arm could go further than a Rubber Hand illusion since, in this case, a limb is attached and included rather than replaced as RHI does.

In line with the discussion above, the visual processing plays a vital role in the task load. Despite the *Hands* condition was kept as simple as possible, it could not be possible to maintain a low cognitive load like in *Graz*. In effect, the processing of visual animation is higher than arrows and fixation cross, showing how the visual processing plays a vital role in the task load. However, the benefits presented by this feedback are reflected in the enhancement of the ERD/ERS signals that consequently produces an improvement in the classification. Supernumerary MI-BCI systems are prominent and possible uses should be explored, especially for VR applications, where customized avatars could be controlled using imaginary non-body signals. In effect, Abdi et al. [49] provide evidences about the usefulness and preferences of having three hands in the execution of some activities (i.e. catching objects).

Additionally, and in line with the previous findings done by Skola and Liarnokapis [34], the embodied training improves the classification performance as well as it elicits stronger and consistent ERS/ERD patterns than the traditional Graz paradigm. However, such comparison, unlike that by Skola and Liarnokapis, is done in VR, i.e.; both conditions were made in an immersive VR scenario, eliminating the bias that exists when the comparison is made with Graz in a monitor-based presentation.

An interesting approach would be studying the sense of agency and ownership of the virtual third-arm using both questionnaires or galvanic skin response (GSR), as done by Bashford and Mehring Bashford and Mehring [38]. This would provide a wider body of knowledge about the use of a supernumerary BCI system. Besides this, an online experiment is mandatory to validate the initial results as well as studies about the handedness of the third arm using left-handed subjects.

Finally, this work also intends to provide premises regarding the role of mental imagery in the exploration of cognitive processes. If we look at the present work from the perspective of embodied cognition, we can argue that supernumerary BCI systems can allow us to study the human ability for body extrapolation and how the mind can be shaped by these new experiences. A discussion is open towards the use of imaginary limbs as a means to control system, extending the human mind constraints imposed by the body.

## 7. Acknowledgements

This study was partly funded by the Coordenação de Aperfeiçoamento de Pessoal de Nível Superior - Brasil (CAPES) - Finance Code 001, and partly by CNPq. We also acknowledge FAPERGS (project 17/2551-0001192-9) and CNPq-Brazil (project 311353/2017-7) for their financial support. Special thanks are also due to Petrobras for the support through the Annelida re-search project. The first author would like to thank to the SDAS Research Group (www.sdas-group.com) for its valuable support.

## Footnotes

1 In order to show visibly the phenomena, we used the EEG data from the subject who obtained the best classification rates [26].

